# A national ‘safe and just operating space’ for all in India: Past, Present and Futurea

**DOI:** 10.1101/390138

**Authors:** Ajishnu Roy, Kousik Pramanick

## Abstract

With 1.3 billion populaces on the commencement of the 21^st^ century, India is currently impending towards upholding a subtle equilibrium between persisting social development and well-being without depleting existing biophysical resources at the national level or surpassing global average per capita obtainability. In this paper, we have structured a top-down per capita framework to explore national ‘safe and just operating space’ (NSJOS) to apprehend not only past fluctuations that bring about the present conditions but also the plausible future consequences, with India as a case study. Coalescing 27 indicators, all pertaining to Sustainable Development Goals (except – SDG 17), accompanied by their corresponding environmental boundaries or preferred social thresholds, present study probes into both biophysical (for environmental stress) and social development (for social deficit) attributes of India. This analysis shows India has already crossed three of seven dimensions of biophysical boundaries (freshwater, nitrogen and phosphorus use). Also, at the existing rate, India is going to cross the remainder of the boundaries within 2045-2050 (climate change, arable land use, ecological and material footprint). Of 20 indicators used for social development, only five have already or will meet corresponding desired thresholds of United Nations Sustainable Development Goals 2015. Using tendencies of past variations, the results indicate that if lowest per capita consumption can be attained and uphold, even with projected population growth, total consumption of four biophysical resources (climate change, nitrogen use, ecological and material footprint) can be slashed from today’s level in 2050. Adaptations in national policy are indispensable if India wants to accomplish sufficiency in biophysical resources whilst bestowing social equity in access and exploitation of those resources towards the continuance of social developments in forthcoming times.

## Introduction

We now have an increasing understanding of the biophysical processes, that not only regulate the stability of the Earth-system thresholds, but also fuel the advancement of our society. Coupled with that, anthropogenic pressures continue to upsurge on the planet swiftly. Every single nation in this 21^st^ century is going through a phase of interconnected causalities from degradation of the environment, deprivation in society and an ineffective economy. Anthropogenic role in geology and ecology has triggered the onset and rapid progress of a new epoch, the ‘Anthropocene’ (Steffen et al., 2011; Waters et al., 2016). Human deeds have pushed Earth-system into a “less biologically diverse, less forested, much warmer, and probably wetter and stormier state” (Steffen et al., 2007). To tackle this on a global scale, the concept of sustainable development has emerged, which is “development that meets the needs of the present without compromising the ability of future generations to meet their own needs” according to Brundtland Commission Report (1987). In 1992, United Nations Conference on Environment and Development (UNCED, Rio de Janeiro Earth Summit), Agenda 21, calls for sustainable development indicators (SDIs) to “provide solid bases for decision-making at all levels and to contribute to a self-regulating sustainability of integrated environment and development system”. United Nations has set 17 Sustainable Development Goals (SDGs) and 169 targets, emerging from Millennium Development Goals (MDGs) (Sachs, 2012) in 2015. This is the first UN approved framework that all nations have agreed towards a ‘broad and universal policy agenda’ that addresses environmental stewardship, human social deprivations and economic equity in an integrated way (UN General Assembly, 2015). These SDGs incorporate all three columns of sustainable development, i.e. environmental goals (climate action, life below water, life on land etc.), social goals (zero hunger, no poverty, gender equality, peace and justice and strong institutions etc.) and economic goals (reduced inequalities, decent work and economic growth etc.). Two major approaches have surfaced to track sustainability, (1) planetary boundaries (PBs) and (2) safe and just space (SJS) framework, under the doughnut economy (DE). In 2009, Rockström et al. introduced a new concept of ‘planetary boundaries’ framework to ascertain ecological or environment thresholds and assess level of consumption of nine biophysical resources related to precarious Earth-system processes (climate change, rate of biodiversity loss, nitrogen and phosphorus cycles, stratospheric ozone depletion, ocean acidification, global freshwater use, change in land use, atmospheric aerosol loading and chemical pollution) whose transgressions risk altering the planet’s Holocene-like steady state of the past 11,500 years ago (Rockström et al. 2009a, Rockström et al. 2009b) . Then Steffen et al. (2015) revised this framework (viz. change in biosphere integrity, land-system change, introduction of novel entities) towards a global scale aggregated evaluation of biophysical thresholds and consumption level. In 2012, Raworth devised 11 dimensions of social foundation based on United Nations Conference on Sustainable Development (Rio+20, 2012) (water, income, education, resilience, voice, jobs, energy, social equity, gender equality, health and food). This framework also has been updated in 2017 to 12 dimensions (viz. food, health, education, income and work, peace and justice, political voice, social equity, gender equality, housing, networks, energy, water) (Raworth, 2017a, 2017b). Dearing et al.’s (2014) case study of two Chinese localities (Erhai lake-catchment, Yunnan province and Shucheng County, Anhui province, China) is a bottom-up analysis which defines ecological processes and control variables based on local environmental conditions of study locations. Nykvist et al. (2013) used a top-down approach to realize national shares of four planetary boundaries (climate change, freshwater use, land-system change, and nitrogen) across 61 countries. Cole et al. (2014), using an assortment of both top-down and bottom-up approaches, designated sustainable development in terms of ‘national barometer’ of South Africa that included both planetary boundaries and doughnut economy frameworks. In this work, they had modified some indicators of both frameworks (arable land use, air pollution and marine harvesting under the PB framework; health care, household goods, safety of SJS framework). Hoff et al. (2014) quantified Europe’s footprint using the PB framework. Dao et al. (2015) applied the PB framework to analyse the sustainability of Switzerland. Kahiluoto et al. (2015) assessed nitrogen and phosphorus boundaries of Ethiopia and Finland. Carpenter and Bennett (2011) have worked on improvising planetary boundary of phosphorus. Recently O’Neill et al. (2018) have downscaled these two frameworks to the nation-scale analysis of 150 nations accompanied with the ushering of new indicators (e.g. eHANPP, ecological footprint, material footprint, life satisfaction, healthy life expectancy, nutrition, social support, democratic quality etc.). More recently, Dao et al. (2018) have analysed the environmental limits of Switzerland in accordance with global limits based on the PB framework. They have analysed PBs related to climate change, ocean acidification, nitrogen and phosphorus loss, land cover anthropisation and biodiversity loss. Though they have used consumption-based indicators in their study, specific socio-economic developmental indicators are absent in their work. In this work, we have tried to unearth answers to the following: (1) How can we downscale both PBs and SJS framework to a national scale more precisely? (2) How can we comprehend changes in dimensions of PBs and SJS with time in order to contextualise their contemporary values? (3) How can we utilize the past trends in biophysical consumptions (PBs) to project probable future consequences at a national scale? (4) How can we understand the interactions or connections among the dimensions of PBs and SJS? (5) How can we summarise and communicate SDG progress in such a way that focus national accomplishments and primaries? Our analysis measures the national performance of India on 28 dimensions, comprising both PBS and SJS frameworks, and provides important outcomes of the relationship concerning biophysical resource use and well-being for India. Our work has been explained herein few steps, (1) we present our methodology and results of our case study on India, (3) we explore interlinks between dimensions of PBs and SJS, (4) we project probable future scenario of biophysical consumption for India, (5) we discuss applicability of PB-SJS framework as a tool in policymaking with local-regional-global links and (6) finally, we discuss limitations of our study and scopes and provisions of further research improvements. In a simple way, we have tried to understand how close India to its ‘safe’ environmental boundaries are (i.e. national biophysical ceiling) (for climate change, freshwater use, arable land use, nitrogen use, phosphorus use, ecological and material footprint) and what proportion of the population lives below ‘just’ social floor (i.e. national social foundation) (for education, energy, food, gender equality, health, housing, income and work, networks, peace and justice, political voice, social equity, water and sanitation). This study is to be used as a study of cautionary warning that exposes the risks that might deter India’s ability to meet its national sustainable development goals as per UN SDG 2015 standard.

## Data and Method

### a. Biophysical Indicators

Though we have mostly adopted Rockström et al.’s (2009b) and Steffen et al.’s (2015) approach of planetary boundaries framework, we have adjusted all of the indicators and boundaries to ensemble national scale and circumstances of India. We have used five indicators as per updated planetary boundaries framework of Steffen et al. (2015) (climate change, nitrogen flow, phosphorus flow, land-system change and freshwater use) and two of O’Neill et al.’s (2018) (ecological and material footprint).

***Climate change***: Rockström et al. (2009b) have calculated climate change boundary based on global ‘atmospheric carbon dioxide concentration (parts per million by volume)’ and ‘change in radiative forcing i.e. energy imbalance at top-of-atmosphere (W m^-2^)’. Cole et al. (2014) and O’Neill et al. (2018) have used ‘annual direct CO2 emissions (Mt CO2)’ and annual per capita CO_2_ emission (t CO_2_), respectively. We have measured climate change in terms of GHG emission per capita per year. According to Emissions Gap Report (UNEP, November 2017), ‘emissions of all greenhouse gases should not exceed 42 GtCO2-e in 2030 if the *2’A* target is to be attained with higher than 66 per cent chance.’ Hence, we have divided 42 GtCO2-e with world population to get per capita global scale boundary of 5.75t CO2-e year^-1^ (2014).

***Freshwater use***: Rockström et al. (2009b) have estimated planetary boundary of freshwater use is the maximum withdrawal of 4000 km^3^ y^-1^ blue water from rivers, lakes, reservoirs, and renewable groundwater stores. Steffen et al. (2015) and O’Neill et al. (2018) have followed this estimate, while Cole et al. (2014) have used annual consumption of available freshwater resources (Mm^3^ per year) We have divided the most accepted value of 4000 km^3^ y^-1^ water with world population to get per capita global scale boundary of 574.86 km^3^ y^-1^ (2010).

***Arable land use***: According to Rockström et al. (2009b), the planetary boundary of land use is less than 15% of global ice-free land cover converted to cropland per year (which is 1995 Mha). Steffen et al. (2015) have measured this in terms of ‘area of forested land as % of original forest cover’ and advocated to maintain a minimum of 75% of global original forest cover (for tropical, temperate and boreal 85%, 50% and 85%, respectively). Cole et al. (2014) have used ‘rain-fed arable land converted to cropland (%)’. O’Neill et al. (2018) have used ‘embodied human appropriation of net primary productivity (eHANPP)’ (t C per capita per year). We have divided globally available and safely maximum usable land of 1995 Mha with world population to get per capita global scale land use boundary of 0.27ha year^-1^ (2015).

***Nitrogen use*** Rockström et al. (2009b) have measured this boundary in terms of ‘amount of N_2_ removed from the atmosphere for human use (millions of tonnes y^-1^)’ (which was 35 million tonnes y^-1^). According to Steffen et al. (2015), the planetary boundary of global nitrogen flow is 62 Tg N y^-^1 from industrial and intentional biological fixation. O’Neill et al. (2018) have followed Steffen et al.’s method for this. Cole et al. (2014) have used the nitrogen application rate of maize production (kg N ha^-1^). We have divided 62 Tg N y^-1^ with world population to get per capita global scale boundary of 8.4kg N year^-1^ per capita (2015). *Phosphorus use:* Rockström et al. (2009b) have measured this boundary in terms of ‘quantity of phosphorus flowing into the ocean (millions of tonnes y^-1^)’ (which gave global boundary of 11 million tonnes y^-1^). But, according to Steffen et al. (2015), the planetary boundary of global phosphorus flow is 6.2 Tg N y^-1^ mined and applied to erodible (agricultural) soils. O’Neill et al. (2018) have followed the same method as Steffen et al.’s for this. Cole et al. (2014) have measured it in terms of ‘total phosphorus concentration in dams (mg/L)’. We have divided 6.2 Tg N y^-1^ with world population to get per capita global scale boundary of 0.84kg P year^-1^ (2015).

***Ecological footprint (EF)***: This is used to measure how much biologically productive land and sea area a population requires to produce the biotic resources it consumes as well as absorb the CO2 emissions it generates, using prevailing technology and resource management practices (Borucke et al., 2013). This is an aggregation of six components (cropland, forest land, fishing grounds, grazing land, built-up land, and carbon land), and can be compared to biocapacity (i.e. total available area of biologically productive land and sea area). O’Neill et al. (2018) first used this ecological footprint in the context of the planetary boundaries framework. According to Global Footprint Network, 12 billion ha biologically productive land and sea area is available in the world. We have divided 12 billion ha with world population to get per capita global scale boundary of 1.66gha year^-1^ (2013).

***Material footprint (MF)***: According to Wiedmann et al. (2015), it (also known as raw material consumption, RMC), measures the amount of used material extraction (minerals, fossil fuels, and biomass) associated with the final demand for goods and services, irrespective of the location of the extraction. It includes the embodied raw materials related to imports and exports and is, therefore, a fully consumption-based measure. The global material footprint has been estimated at 70 Gt y^-1^ (i.e. 10.5 ton per capita in 2008, by Wiedmann et al. 2015), and it was capped to 8 ton per capita has been suggested as a sustainable level, by Dittrich et al. (2012). According to Dittrich et al. (2012), global material extraction should not exceed ~50 Gt y^-1^, based on the material used in 2000 (50.8 Gt). We have divided 50 Gt y^-1^ with world population to get per capita global scale boundary of 7.18t year^-1^ (2010).

These dimensions along with their respective indicators, boundary, current status, data source and boundary crossing time (at BAU rate) are explained in Table 1.

**Table 1:** Dimensions and indicators of the ecological ceiling (Planetary boundaries concept) for India

| No. | Dimension (related SDG) | Indicator | Boundary (global, per capita) | Year | Current Status | Change since initial year (+ increase, - decrease) | Data Source | Boundary crossing time (BAU) |
| --- | --- | --- | --- | --- | --- | --- | --- | --- |
| 1 | Climate change (SDG 13) | GHG emissions including Land-Use Change and Forestry (t CO<sub>2</sub>-e) Per Capita Per Year | 5.75t CO<sub>2</sub>-e year<sup>-1</sup> (2014) | 1990 - 2014 | 2.47 t CO<sub>2</sub>-e (2014) | +46.95 % | Climate Analysis Indicators Tool (CAIT) | 2045-2047 |
| 2 | Freshwater use (SDG 6) | Total water withdrawal (m<sup>3</sup>) Per Capita Per Year | 574.86m<sup>3</sup> year<sup>-1</sup> (2010) | 1973 - 2012 | 602.3 m<sup>3</sup> (2010) | +3.07% | Aquastat | 2007-2008 |
| 3 | Arable land use (SDG 15) | Agricultural land area (ha) Per Capita Per Year | 0.27ha year<sup>-1</sup> (2015) | 1961 - 2015 | 0.13 ha (2015) | -64.61% | FAOSTAT, World Bank | 2035 |
| 4 | Nitrogen use (SDG 14) | Nutrient nitrogen N use (kg) Per Capita Per Year | 8.4kg N year<sup>-1</sup> (2015) | 2002 - 2015 | 14.07 kg N (2015) | +27.51 % | FAOSTAT | 2002 |
| 5 | Phosphorus use (SDG 14) | Nutrient phosphate P<sub>2</sub>O<sub>5</sub> (kg) Per Capita Per Year | 0.84kg P year<sup>-1</sup> (2015) | 2002 - 2015 | 5.65 kg P (2015) | +30.54 % | FAOSTAT | Before 2002 |
| 6 | Ecological footprint | Ecological footprint | 1.66gha year<sup>-1</sup> | 1961 - | 1.06 gha | +43.87 % | Global Footprint | 2020-2025 |

|  | (EF)<br>(SDG 14,<br>15) | (gha) Per<br>Capita Per<br>Year | (2013) | 2013 | (2013) |  | Network<br>(GFN) |  |
| --- | --- | --- | --- | --- | --- | --- | --- | --- |
| 7 | Material<br>footprint<br>(MF)<br>(SDG 12) | Material<br>footprint<br>(t) Per<br>Capita Per<br>Year | 7.18t<br>year<sup>-1</sup><br>(2010) | 2000<br>-<br>2010 | 3.56 t<br>(2010) | +30.61<br>% | UNData<br>(UNSD) | 2025-2030 |

Boundaries, that have not yet been crossed, are shown in green and yellow represents boundaries that have been crossed. Boundaries that have been crossed and reached a critical level that deserves serious attention are represented in red.

### b. Social development Indicators

We have followed framework consisting 12 dimensions of Raworth (2017a).

***Education***: SDG 4 targets on ensuring inclusive and equitable quality education and promotion of lifelong learning opportunities for all. We have chosen 3 indicators to understand primary, secondary and adult education (adult literacy rate, children remained in school (of primary school age) and secondary school enrolment) to reflect achievements and outcomes across diverse population age groups. We have focused on these for three reasons. First, primary education is the very basic one should have, especially to be able to cope up with a growing economy (India). Second, without receiving a comprehensive subject- or skill-oriented knowledge during secondary school years, the young population might become ill-equipped for tertiary education or the workforce, also can be engaged to activities with negative effects on social well-being (as - radicalization by militants, unplanned teenage pregnancy, juvenile crimes etc.). Third, education paradigm was not even sufficiently developed in India a few decades ago for most of the people (i.e. lack of access to educational institutions, education-aiding equipment and technology etc.). Therefore, monitoring adult literacy becomes important for both themselves and their next generations. We have used the data from the World Bank’s World Development Indicators (WDI). Similar to the other percentage indicators, a threshold of 10% or less was chosen for children out of the school of primary school age and 90% or more was chosen for secondary school enrolment and adult literacy rate as - universal access to education does not imply 100% enrolment, especially for countries like India.

***Energy***: SDG 7 focuses on ensuring access to affordable, reliable, sustainable and clean energy for all. About 1 billion people currently do not have access to electricity. 3 billion people rely on polluting fuel (like – fuelwood, charcoal, crop residue, animal dung, dry leaves) to cook food, which in turn resulting in 4 million premature deaths per year, mostly among women and children, that are due to household air pollution (SDG 7 tracking report, 2018) . Our assessment of deprivations in access to energy includes both electricity and the quality of (clean) cooking facilities. We have measured energy using two indicators, (1) ‘access to electricity (% of populations)’ and (2) ‘access to clean fuels and technologies for cooking (% of the population)’, obtained from the World Bank’s WDI. The threshold for energy was set at 90% or more for both indicators.

***Food***: The target of SDG 2 is ending hunger, achieving food security and improved nutrition for all. We have measured social development related to food using two indicators, (1) ‘prevalence of undernourishment (% of the population)’, and (2) ‘average calorific intake of food & drink (kcal/capita/day)’. The first indicator, from the World Bank’s WDI, is selected keeping in mind the occurrence of undernourishment and malnutrition in almost all of the developing countries, like – India. The second indicator (by UN FAO) is an average calorific intake of food and drink, with unit - kilocalories (kcal) per capita per day. The physiological requirements for an average adult remain between 2100 and 2900 kcal per day (for average women and men during moderate physical activity). However, this calorific requirement range exceeds for individuals associated with heavy manual labour or athletic activity (Smil, V., 2000). We have used 2700 kcal or more per capita day y^-1^ as the desired threshold.

***Gender Equality***: The focus of SDG 5 is achieving gender equality via empowering all women. It would be ideal to assess the extent of gender inequality to understand women and men’s roles and status in political and economic life. We have measured this using one indicator - ‘proportion of seats held by women in national parliaments (%)’ from the World Bank’s World Development Indicators. The indicator value is calculated such that if women held exactly half of all parliamentary seats (i.e. 50%), that should be non-biased to both genders. Thus, achieving 50% seats in parliament has been taken as the desired threshold.

***Health***: Ensuring healthy lives and promoting well-being for all at all ages is the focus of SDG 3. We have used two indicators to assess shortfalls in access to health care in India: (1) ‘life expectancy at birth, total (years)’ and (2) ‘mortality rate, <5 years (per 1,000 live births)’ from the World Bank’s WDI, both selected for being recognized proxies for wider health outcomes. First indicator, life expectancy at birth indicates the number of years a newborn infant would live if prevailing patterns of mortality at the time of its birth were to stay the same throughout its life. According to the Human Development Report (UNDP 2015), 70 years or more life expectancy at birth is selected here as a desirable threshold. The second indicator, under-five year mortality rate is the probability per 1,000 that a newborn baby will die before reaching age five, based on age-specific mortality rates of the specified year. According to WHO (2015), the international target for all countries to reduce under-five years age mortality to at least as low as 25 per 1,000 live births by 2030. Thus, 25 or less per 1000 live births has been set as the desired threshold here.

***Housing***: SDG 11 focus on making cities and human settlements inclusive, safe, resilient and sustainable. We have measured it with ‘population living in slums (% of urban population)’ from WDI (World Bank). Slum housing is defined as having at least one of the following four characteristics - lack of access to improved drinking water, lack of access to improved sanitation, overcrowding (>3 persons per room) and dwellings made of non-durable material. As most of the Indian people presently live in rural areas, an indicator that measures deprivations in conditions in rural houses of India would have been more appropriate, along with the used indicator. We have set the threshold at 10% or less of urban population living in slums.

***Income and Work***: SDG 1 focus on ending poverty in all its forms everywhere. Promoting sustained, inclusive, sustainable economic growth full and productive employment and decent work for all is the goal of SDG 8. We have measured income with (1) ‘poverty headcount ratio at $1.90 a day (2011 PPP) (% of population)’ and work with (2) ‘unemployment, youth total (% of total labour force, 15-24 years)’ both from WDI (World Bank). The first indicator is defined as the poverty threshold at $1.90 a day using 2011 international prices. Although the goal is having 100% of the population living above the $1.90 a day line, we have used a threshold value of 95% in our analysis. The second indicator is youth unemployment which measures the proportion of young people (aged 15-24 years) who are seeking but unable to find work (International Labour Organization, ILO estimation). The unemployment rate means the share of the labour force that is without work but available for and seeking employment. We have used 94% or more people are employed, i.e. 6% or less unemployed people below this line is the desired threshold for this indicator.

***Networks***: SDG 9 focus on building resilient infrastructure, promoting inclusive and sustainable industrialization and fostering innovation. Under this goal, target 9.c. focus on significantly increasing access to information and communications technology and strive to provide universal and affordable access to the Internet in the least developed countries. Network was measured using ‘individuals using the internet (% of the population)’ provided by WDI of the World Bank. Digital communications networks are important means of generating opportunity, building community and increasing resilience. We have set here 90% or more of the population have access to the internet as the desired threshold for this indicator.

***Peace & Justice***: UN SDG 16 focus on promoting peaceful and inclusive societies for sustainable development, provide access to justice for all and build effective, accountable and inclusive institutions at all levels. We have used two indicators to assess shortfalls in justice and peace, (1) corruption perceptions index (CPI), (provided by Transparency International) and (2) ‘intentional homicides (per 100,000 people)’ (from WDI) respectively. The first indicator, corruption perception index scores countries according to how corrupt their public sector is perceived to be, on a scale of 0 to 10 (up to 2011) and 0 to 100 (2012 onwards) (i.e. highly corrupt to very clean). We have set the desired threshold of 5 or less (up to 2011) and 50 or less (2012 onwards). The second indicator defines the rate of intentional homicide as unlawful death purposefully inflicted on a person by another person. The threshold is set at 100 or fewer homicide deaths per 100,000 population per year.

***Political Voice***: Under SDG 16, target 16.7 aims for ensuring responsive, inclusive, participatory and representative decision-making at all levels. We have measured political voice using voice & accountability index (VAI), provided as a component of the World Bank’s World Governance Indicators (WGI). This index is scored on a scale of 0 to 1 (i.e. very poor performance to very high performance) and includes measures of democracy, vested interests, accountability of public officials, human rights, and freedom of association. The threshold is set at 0.5 or less on this indicator.

***Social Equity***: SDG 10 focus on reducing inequality within and among the countries. The shortfall of social equity is measured with national income inequalities. We have measured social equity using the Gini coefficient provided by the World Income Inequality Database 3.4 (WIID 3.4). Evidence for high-income countries suggests that more equal societies have fewer health and social problems than less equal ones. The threshold was chosen of 70 of 0-100 scale of Gini index of 0.30.

***Water & Sanitation***: SDG 6 focus on ensuring availability and sustainable management of water and sanitation for all. Deprivations in access to water and sanitation services are assessed on the basis of two widely used indicators, (1) ‘improved sanitation facilities (% of the population with access)’ and (2) ‘improved water source (% of the population with access)’ from the World Bank’s WDI. The sanitation indicator measures the percentage of the population using improved sanitation facilities. Although it is preferable that 100% of the population should have access to improved sanitation facilities, we have chosen a threshold of 90% for this indicator in recognition of the fact that most of the Indian population are located in rural areas. Inadequate access to water denotes the proportion of people who do not have access to an improved drinking water source, like - piped household water, public taps, protected wells and springs, or collected rainwater etc. We have set the threshold for this indicator as 90% or more people in India have access (i.e. 10 or fewer people do not have access) to improved water source.

These dimensions along with their respective indicators, threshold, current status, data source and threshold meeting time (at BAU rate) are explained in Table 2.

**Table 2:**
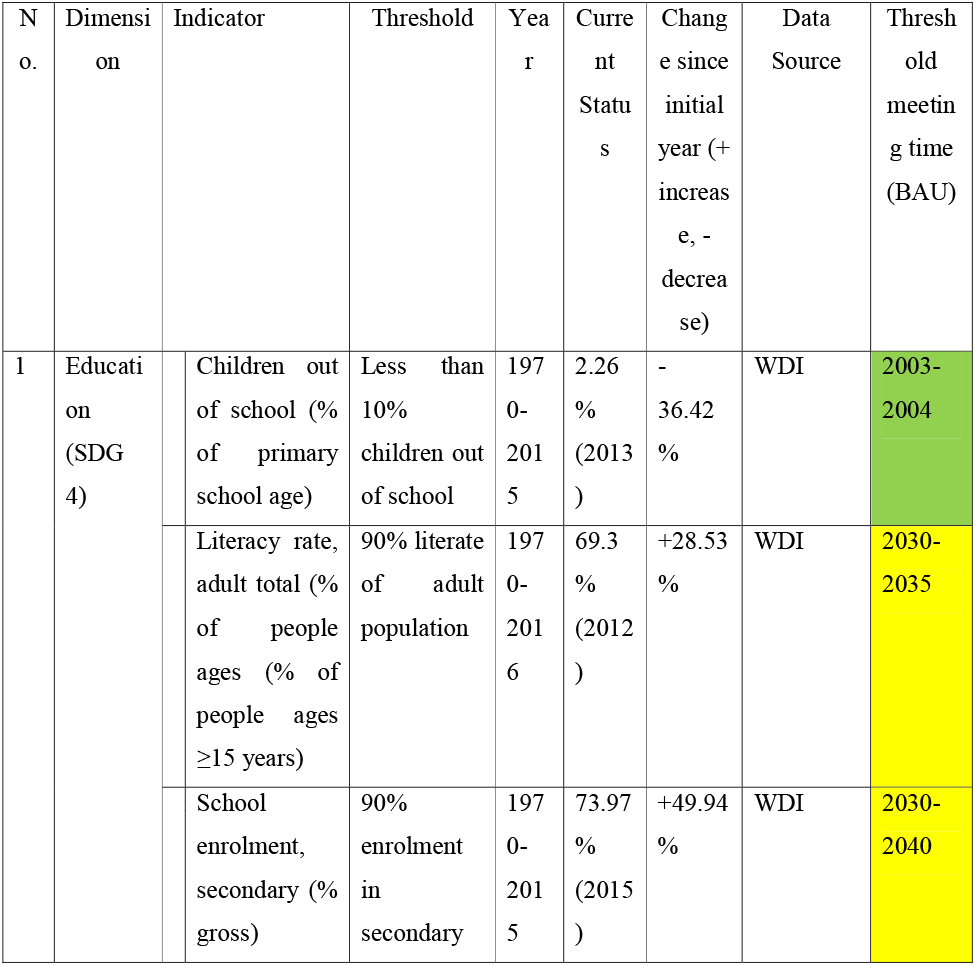
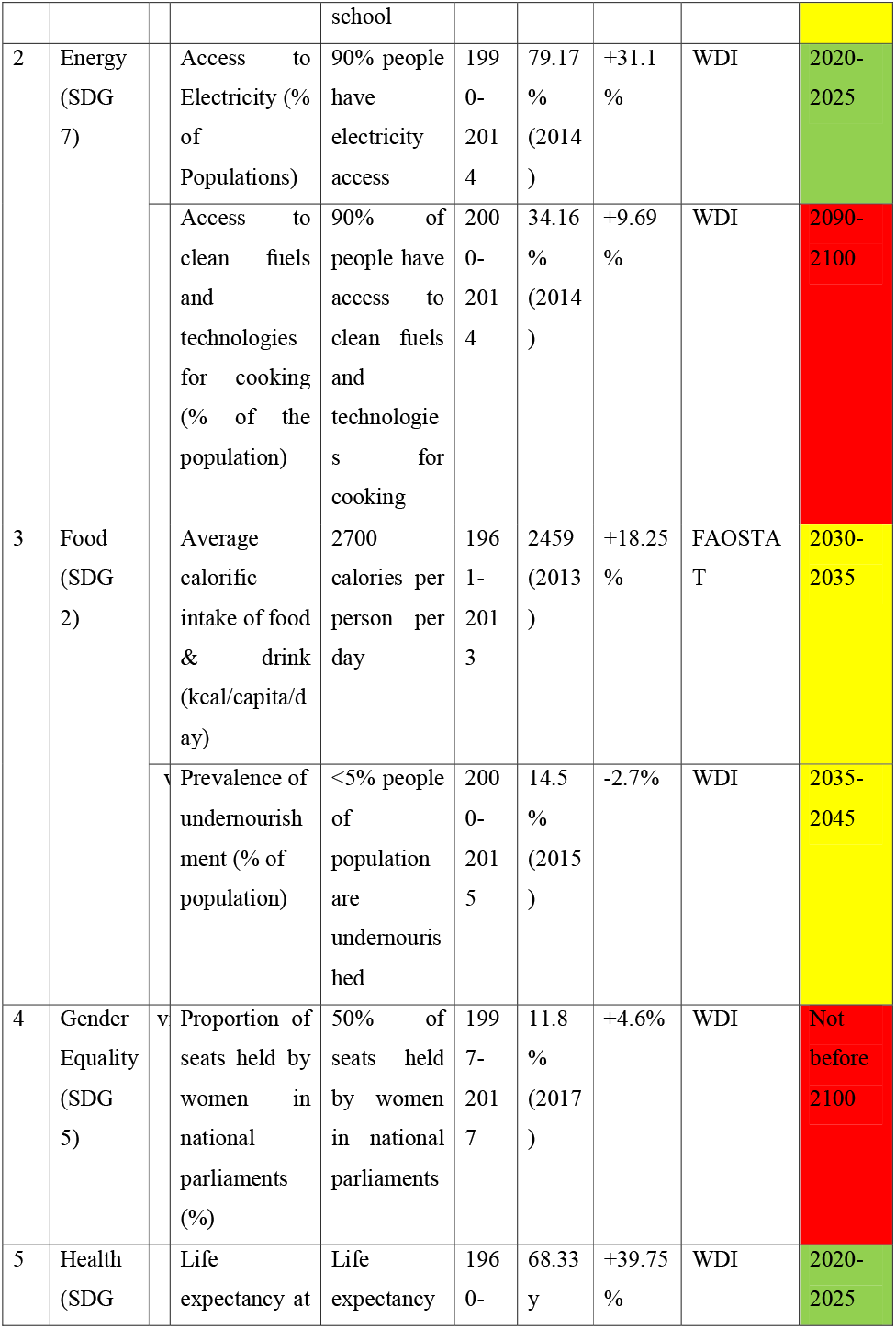
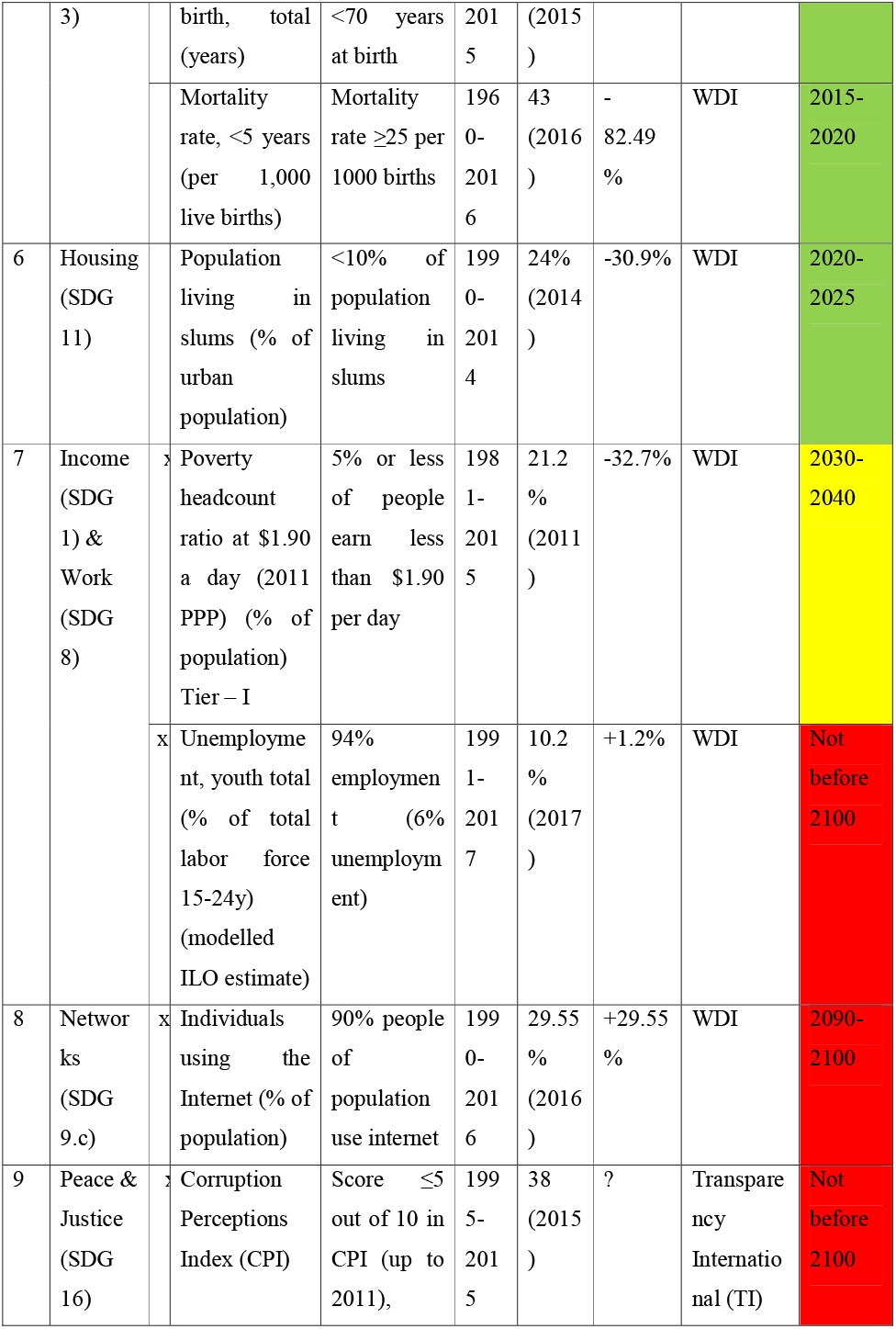
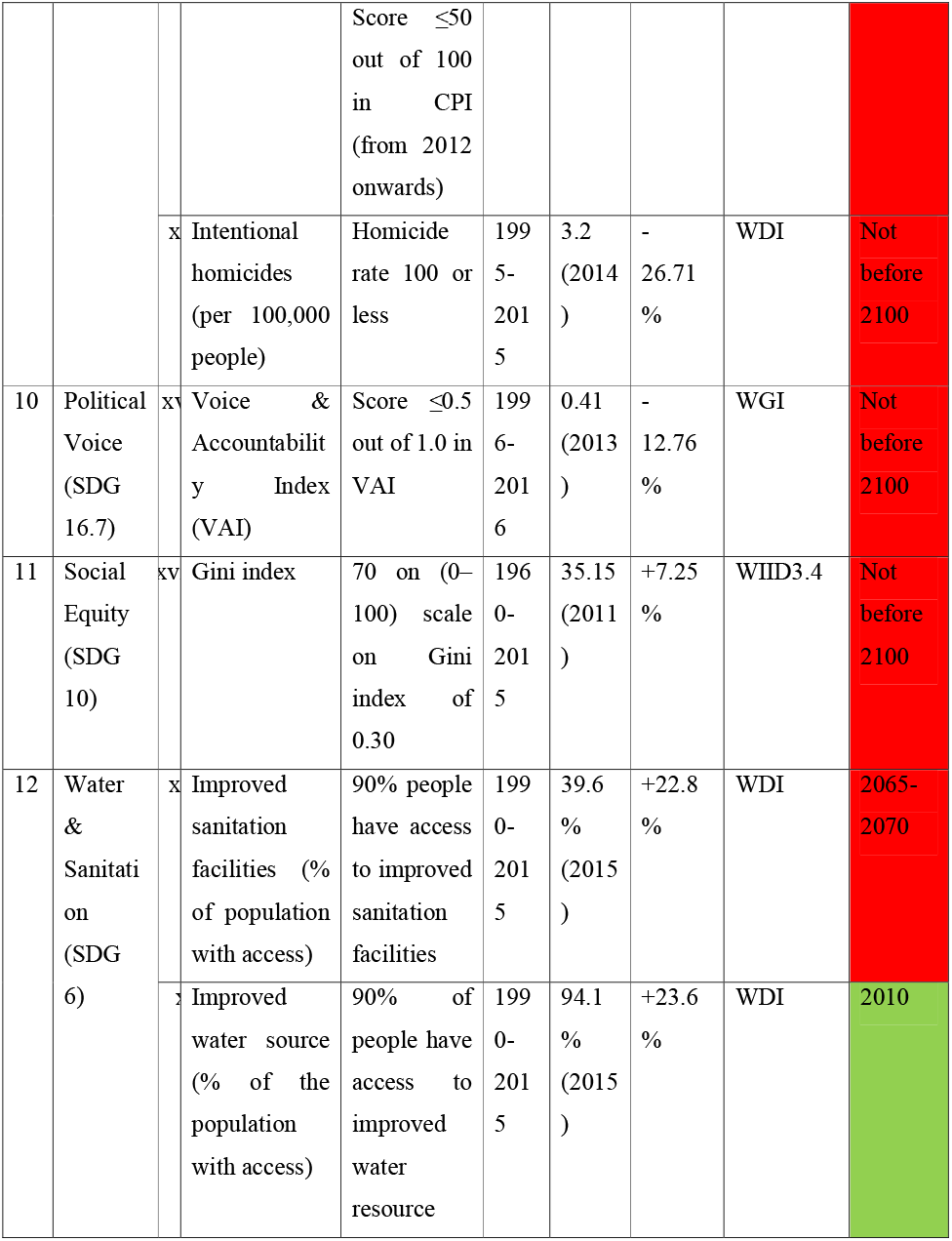
Dimensions and indicators of social foundation (Safe and Just space, SJS of Doughnut economy concept) for India.

Green represents indicators that are going to meet or have already met threshold within UN SGD target time (2030). Indicators that are going meet a few years after that time are shown in yellow and which are going to meet the desired threshold many years after 2030 are shown in red.

To establish, causation between each of biophysical indicators on each of social development indicators and to have a better overview on the associative nature of each pair, we conducted an OLS regression with biophysical indicators as independent variables and each of social development indicators as the dependent variable (Supplementary Table).

### c. Future scenario

As we have calculated all the biophysical indicators on per capita basis, it is possible to project probable future scenario of total consumption. We collected future population projection (2015-2050) data (median range prediction value of 50%) of India from UN DESA (2017 Revision) and then multiplied it with per capita consumption of dimensions of PBs. We have calculated three projection series for each dimension of PBs, (1) with the lowest value that has happened in past year, (2) highest value that has happened in past year and (3) business-as-usual scenario with latest available data.

## Results

### a. Biophysical Indicators

In India, GHG emission, freshwater use, nitrogen use, phosphorus use, ecological footprint and material footprint have increased 46.95% (from 1990), 3.07% (from 1973), 27.51% (from 2002), 30.54% (from 2002), 43.87% (from 1961) and 30.61% (from 2000), respectively. On the other hand, arable land use has decreased 64.61% (from 1961) in India. India has already crossed three of seven dimensions of per capita biophysical boundaries (freshwater use – 2007-2008, nitrogen use – 2002, and phosphorus use – before 2002). If everything remains unchanged, i.e. BAU scenario, India would cross the rest of four dimensions of PB within 2045 (climate change – 2045-2047, arable land use – 2035, ecological footprint – 2020-2025, material footprint – 2025-2030). Three PBs have exceeded their boundaries by 4.5% (freshwater use), 40.3% (nitrogen use) and 85.1% (phosphorus use). Remaining four PBs are within 42.9% (climate change), 51.85% (arable land use), 36.1% (ecological footprint) and 50.5% (material footprint) of exceeding their boundaries (Fig. 1).

**Fig. 1.**
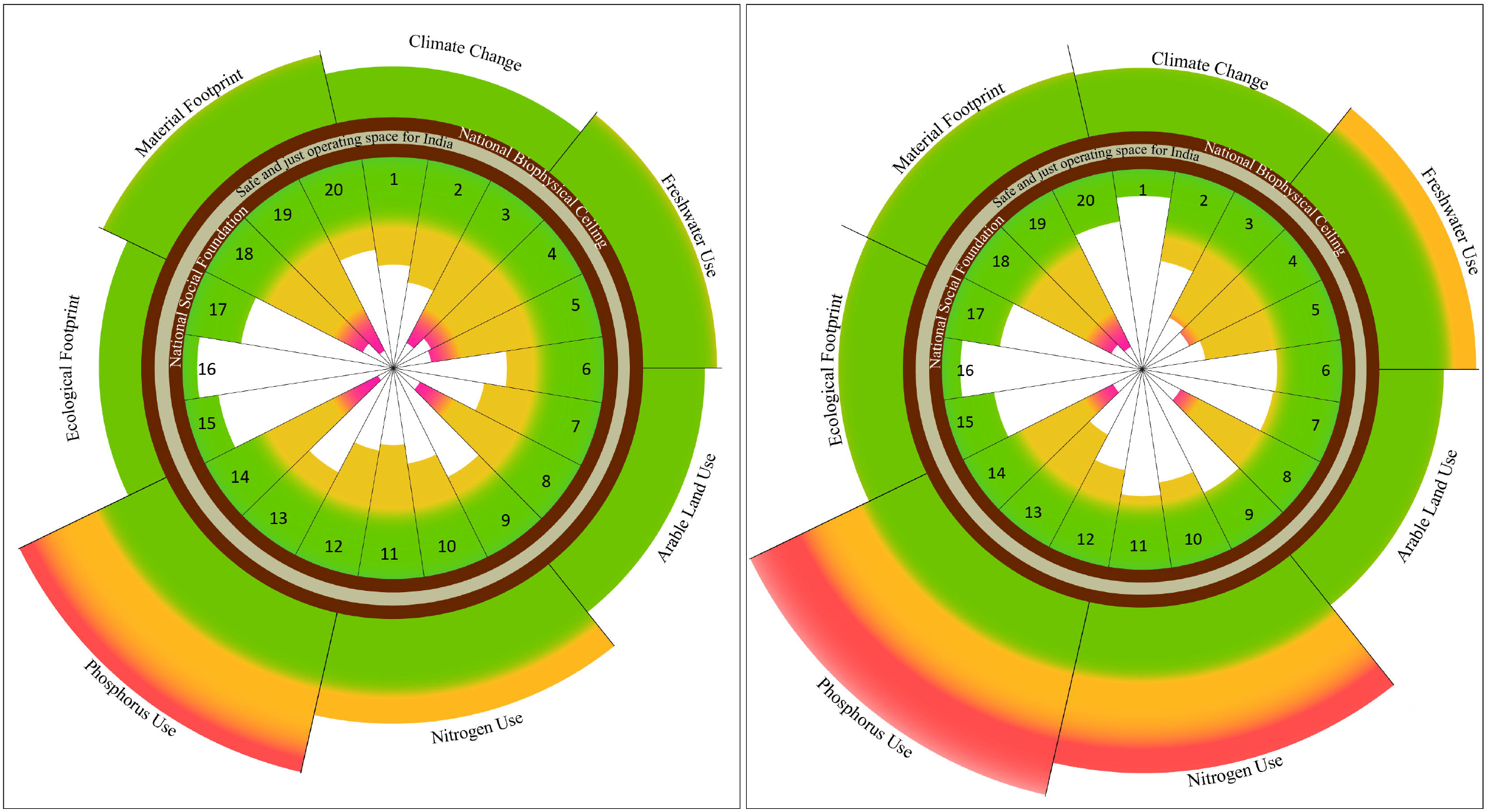
Trends in national barometer for sustainable development in India.

Seven national dimensions of biophysical stress established over national biophysical ceiling outwardly projected and twenty indicators composing twelve dimensions of social deprivation established under national social foundation inwardly projected for India. A and B represent the status of sustainable development of India in 2000 and 2011 respectively. Biophysical indicators are climate change, freshwater use, arable land use, nitrogen use, phosphorus use, ecological footprint and material footprint. Indicators of social development are - (1) children out of school of primary school age, (2) adult literacy rate, (3) secondary school enrolment, (4) access to electricity, (5) access to clean fuels and technologies for cooking, (6) average calorific intake of food and drink, (7) undernourishment, (8) proportion of seats held by women in national parliaments, (9) life expectancy at birth, (10) mortality rate under 5 years, (11) urban population living in slums, (12) poverty headcount ratio at $1.90 a day, (13) youth unemployment, (14) individuals using the internet, (15) corruption perception index, (16) intentional homicides, (17) voice and accountability index, (18) Gini index, (19) improved sanitation facilities and (20) improved water source. Twelve dimensions of the social foundation are education 1-3, energy 4-5, food 6-7, gender equality 8, health 9-10, housing 11, income and work 12-13, networks 14, peace and justice 15-16, political voice 17, social equity 18, water and sanitation 19-20. Green indicates safe operating space for biophysical indicators and thresholds for indicators of social development. Yellow indicates the zone of increasing impact for biophysical indicators and zone of increasing deprivation for indicators of social development. Red indicates the zone of high risk of serious impact for biophysical indicators and zone of the high level of deprivation for indicators of social development. The area between the chocolate rings is the safe and just operating space for sustainable development in India.

### b. Social development Indicators

In India, primary school age children out of school, undernourished population, mortality rate in less than 5y age children (per 1000 live births), urban slum living population, poverty headcount ratio, intentional homicides (per 1,00,000 people), voice and accountability index score have decreased 36.42% (from 1970), 2.7% (from 2000), 82.49% (from 1960), 30.9% (from 1990), 32.7% (from 1981), 26.71% (from 1995) and 12.76% (from 1996), respectively. On the other hand, adult literacy rate, secondary school enrolment, access to electricity, access to clean fuels and technologies for cooking, average calorific intake of food & drink, seats held by women in national parliament, life expectancy at birth, youth unemployment, internet using population, social equity, improved sanitation facilities using population and improved water source availing population have increased 28.53% (from 1970), 49.94% (from 1970), 31.1% (from 1990), 9.69% (from 2000), 18.25% (from 1961), 4.6% (from 1997), 39.75% (from 1960), 1.2% (from 1991), 29.55% (from 1990), 7.25% (from 1960), 22.8% (from 1990) and 23.6% (from 1990), respectively. For the 12 dimensions of SJS framework, only one (peace and justice) has been non-deprived. Of 20 indicators that we analysed for India, only five were not socially deprived, which are – ‘children out of school (% of primary school age)’, ‘CPI’, ‘intentional homicides (per 100,000 people)’, ‘VAI’ and ‘improved water source (% of population with access)’. However, among the rest 15 indicators, only one is getting more distant from the threshold, namely youth unemployment of 15-24y; all the remaining 14 indicators were coming are closing in towards their respective thresholds. Five of the 14 indicators are showing much improvement over the years (Secondary school enrolment, access to electricity, life expectancy at birth, Mortality rate of <5 years and urban population living in slums) and in a BAU scenario, they might reach their thresholds within a few years. Most deprivation exists for eight dimensions – energy, gender equality, income and work, networks, peace and justice, political voice, social equity, water and sanitation. Least deprivation exists for three dimensions - education, health, housing.

Thus, it is clear that exceeding the environmental safe limits have serious consequences for the national security of energy, food, water, job and health; which in turn, potentially, might affect the national economy and international trades. So, it is evident that national policy decisions on socio-economic development should take environmental costs into account if these need to be sustainable.

Trends of changes in both biophysical and social development indicators, over time, are shown in Fig. 2.

**Fig. 2.**
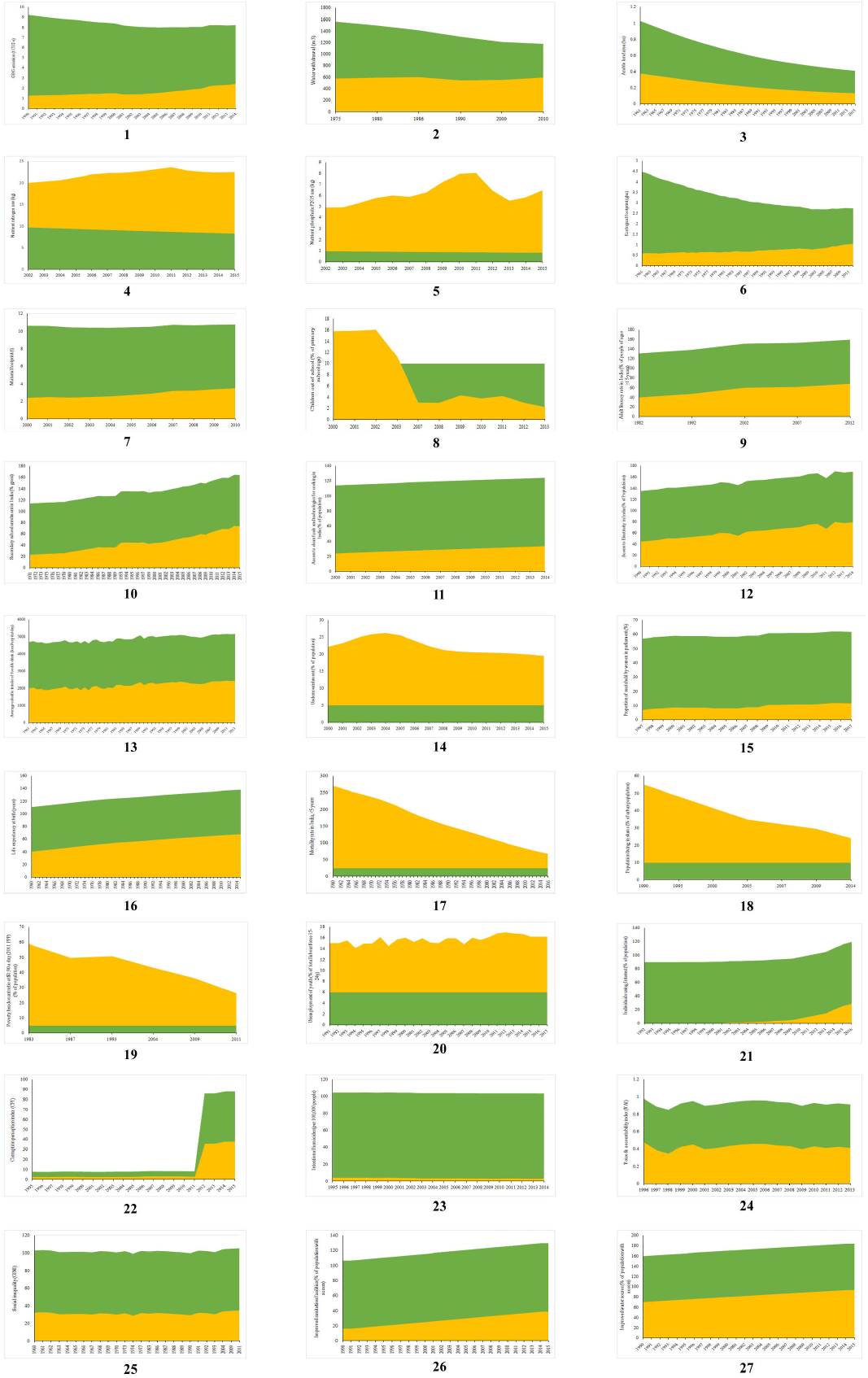
Changes in per capita biophysical indicators (1-7) and indicators of social development (8-27) of sustainable development in India with time.

Green indicates global per capita boundaries for biophysical indicators and thresholds for indicators of social development. Yellow indicates values of indicators for India.

Biophysical indicators are – (1) GHG emission, (2) water withdrawal, (3) arable land, (4) nitrogen use, (5) phosphorus use, (6) ecological footprint and (7) material footprint. Indicators for social development are - (8) children out of school of primary school age, (9) adult literacy rate, (10) secondary school enrolment, (11) access to electricity, (12) access to clean fuels and technologies for cooking, (13) average calorific intake of food and drink, (14) undernourishment, (15) proportion of seats held by women in national parliaments, (16) life expectancy at birth, (17) mortality rate under 5 years, (18) urban population living in slums, (19) poverty headcount ratio at $1.90 a day, (20) youth unemployment, (21) individuals using the internet, (22) corruption perception index, (23) intentional homicides, (24) voice and accountability index, (25) Gini index, (26) improved sanitation facilities and (27) improved water source.

### c. A ‘safe and just’ India in 2050

If we can cap GHG emission at lowest per capita level (i.e. 1990 level) of India, even with grown population projection level of 2050, GHG emission can be lowered 31.98%. But at the present rate (which is also the highest per capita rate), GHG emission will increase 22% in 2050. At the present rate (2010) of per capita water use, in 2050 it will increase 23.83%. Likewise, at a high rate (1986), it will even increase more (24.62%) in 2050. But, if India can attain the lowest per capita water use rate (1990), even with the 2050 population level, the increase will be lower (16.83%). In 2050, nitrogen use is going to increase by 23% (at BAU rate) or even 26.64% (at highest per capita rate of 2011). But it can be lowered to 5.85% decrease if the lowest per capita rate (of 2002) can be attained in 2050. Phosphorus use is going to increase 31.03% (at BAU rate) or even 45.48% (at highest per capita rate of 2011) in 2050. However, it can be decreased by 0.7% from the present level in 2050 if India can attain the lowest per capita use level (of 2002). At the recent rate of per (2013) capita consumption, ecological footprint is going to increase 22.93% in 2050. But it can be 27.8% decreased from recent level if the lowest per capita level (1965) can be attained. Likewise, at a recent rate of per capita consumption (2010), the material footprint is going to increase 25.79% in 2050. But it can be 6.49% decreased from recent level if the lowest per capita level (2000) can be attained in 2050. Probable consequences of biophysical resource consumptions accompanied with lowest and highest rate per capita consumption for India are shown in Fig. 3.

**Fig. 3.**
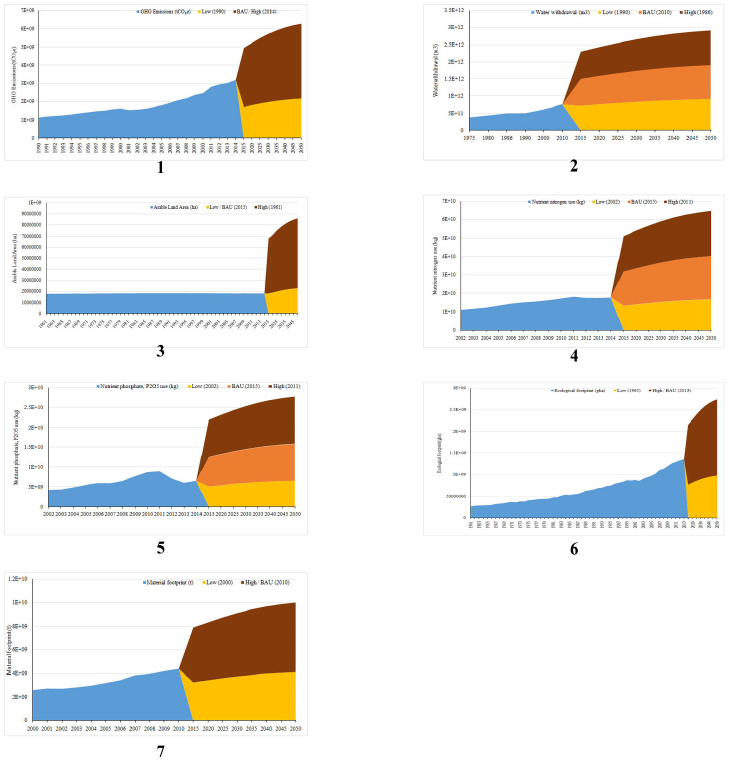
Future scenario of biophysical indicators for India up to 2050.

Blue indicates changes in total values of consumption of biophysical indicators for India. Brown indicates projected total values at the highest rate of per capita consumption; Orange indicates projected total values at business-as-usual (BAU) rate of per capita consumption, Yellow indicates projected total values at the lowest rate of per capita consumption.

## Discussion

Though this field is nascent, there have been a lot of interdisciplinary studies related to either only PB or SJS framework. Sayers and Trebeck (2015a) and Sayers (2015b) have applied DE framework both for Welsh and UK. Chapron et al. (2017) have advised enforcing environmental laws as tools to constraint human impacts on the environment through staying under safe planetary boundaries. There have been a lot of debate surrounding a suitable indicator for ‘biosphere integrity’ (Samper, 2009; Running, 2012; Mace et al., 2014; Newbold et al., 2016), ‘freshwater use’ (Rockström and Karlberg, 2010; Bogardi et al., 2013; Gerten et al., 2013; Heistermann, 2017) ‘introduction of novel entities’ (Sala and Goralczyk, 2013; Persson et al., 2013; Diamond et al., 2015; Villarrubia-Gómez et al., 2017) accompanied by their respective safe boundaries. Some work has also been done on connecting governance with the planetary boundary along with its policy implications (Bierman, 2012, Galaz et al., 2012a, 2012b; Reischl, 2012). There has been a significant amount of works on establishing and applying PB framework in regional scenario (Dearing et al., 2014; Häyhä et al., 2016; Cole et al., 2017; McLaughlin, 2018). Some important works also have been done to establish the connection of PB framework to the food system and nutrients (Kahiluoto et al., 2014, 2015; Campbell et al., 2017; Conijn et al., 2018). Nash et al. (2017) have prepared a framework to apply this PB framework in the marine context. Recently, there have been some criticisms too of this SJS framework (Montoya et al., 2018a, 2018b).

Two previous studies have incorporated sustainability of India based on PB and SJS framework, Nykvist et al. (2013) and O’Neill et al (2018). Nykvist et al. (2013) have considered four planetary boundaries in their report, namely climate change (tCO_2_ per capita y^1^), nitrogen use (kg N per capita y^-1^), freshwater use (m^3^ per capita y^-1^) and land use (ha per capita). According to them, India did not cross per capita PB of climate change (for 2008, either in territorial or consumptive emission). Although, did not cross nitrogen flow PB (2005-2009 average), freshwater use PB (1996-2005) and land use PB (2005-2009). One problem in this study is that it did not include correlated social dimensions (i.e. DE framework or any other). O’Neill et al. (2018) used seven and eleven indicators for PB and DE analysis, respectively. According to them, India crossed only one boundary, climate change PB (ton CO_2_ per capita y^-1^) and not socially deprived in only one threshold, employment (% of the labour force employed).

The primary aim of this study was to evaluate the applicability of SJS framework at the national level in India. We have tried to maintain the original design and concept of the framework as much as possible while deriving results that are meaningful in the Indian national context.

If all Indians ought to lead a prosperous life within the safe limits of planetary boundaries, then resource utilization processes must be fundamentally restructured to enable basic needs at a lower level of resource consumption that does not significantly transgress planetary boundaries. Resource use could be reduced significantly in India with lowering per capita consumption while achieving a more equitable distribution of access to resources among all the people. To focus on sufficiency in biophysical resource consumption, recognizing the overconsumption is a key point which burdens Indian society with a mix of environmental and socioeconomic problems.

We have downscaled PB framework and applied SJS framework at a national scale, for India for the first time, creating an analysis for inclusive sustainable development for India. This work presents the present state and trajectory of a comprehensive yet manageable set of indicators for environmental and social aspects. This work also highlights India’s closeness to environmental boundaries and the nearness from the abolition of social deprivation as per UN SDG 2015 targets. Thus, it creates a preliminary monitoring and communication tool for the government to integrate environmental and social development issues. This study provides insight into the targets for the proposed global UN Sustainable Development Goals that are nationally relevant.

There are a few recommendations that we have come up during this work: (1) Sub-national level database should be established and carefully updated by the government that are publicly available. Because - globally defined boundaries can fall short for many SDG dimensions, where national resource availability limits and local thresholds are more suitable. (2) Data coverage period should be as long as possible along with monthly or at least daily data. Rather than a barrier, this should be utilized as a prospect for data-poor countries to begin a proficient targeted assemblage of comprehensive key data to address respective national to global challenges. (3) The multinational analysis should be done that can yield a comparative overview of national states. Also, each nation can understand which and where to focus to be able to meet UN SDG criteria. (4) Every indicator should be prepared in such a way that each can explore the national context and establish a connection with international academia. (5) It is necessary to use existing data for a nation (like – India) and refine this PB-SJS framework over time as more data are gathered until UN SDG criteria are accomplished. (6) Appropriate indicator to measure progress for the original Steffen’s (2015) and Raworth’s (2017) framework should be developed. (7) To tackle equity in resource consumption for rich and poor countries per capita boundaries should be integrated with steady-state economics (Daly, 1972, 2008; O’Neill, 2012, 2015; Kosoy et al., 2012; Steffen and Smith, 2013), customized for each of the nation’s economy, to solve of differences in the degree of development and the right to develop. (8) To signify the study, multiple variables should be compared for a certain period of time for groups of countries, especially clustering them based on the general understanding of political economy and geography. (9) Further work is necessary for an approach to analyze policymaking and their implementation gaps of each nation for all of the indicators in the SJS framework. Problems might ascend when any PB is only incompletely addressed in a national policy objective. (10) The importance of considering local environmental problems and threats to local ecological resilience should be emphasized during use of this type of methodologies and results. Though SJS framework was developed to highlight and strengthen understanding and awareness about the planetary consequences of different environmental processes due to anthropogenic pressures, it is not that only the planetary problems are significant. SJS framework is to be used as a complementary to the analysis of local and national socio-ecological problems, which need to be addressed in their own significance regardless of the apparent absence of obvious and readily understandable planetary implications. (11) The PB framework (‘safe’ part) should be analyzed and results with policy adaptations are to be strictly implemented mostly in developed countries where social development has already taken place through eradication of deprivation, whereas DE framework (‘just’ part) should be adopted in less developed countries where social development is apparently more important and need of the hour. (12) When comparing the sustainability performance of countries based on SJS framework, developed countries (like – USA, UK, Germany, France etc.) and countries with rapidly growing economies (like – India, China etc.) are to be specially emphasized. These countries have either higher total or per capita impacts on the environment globally, and hence of bigger responsibility. (13) We recommend that every nation (like – India) should act more proactively and adopt policies according to recommendations of international bodies, like - UN, UNFCC, UNDP, UNEP etc. if the country desires to reduce its sustainability deficit. (14) Boundaries for all the dimensions under the PB framework, especially applicable to a national scale, should be established. (15) SJS framework should be accompanied with systems dynamic analysis of the interrelationships between any of the biophysical social conditions. Till now, it only conveys a basis for judging the relative state of current biophysical viability and societal wellbeing on a global scale. (16) There are some dimensions of biophysical resource consumption related to PB framework that done have any unanimously selected representative indicators along with corresponding boundaries, specially customized to fit national or sub-national level analysis, such as – change in biosphere integrity, stratospheric ozone depletion, ocean acidification, atmospheric aerosol loading and the introduction of novel entities. It should be a priority. (17) It should be kept in mind that just identifying indicators for the PB framework and their monitoring is not enough. If proper checkpoints need to made for these and for policy implications, drivers for each PB dimension for every nation should be identified first. (18) How to plan a future for a nation under safe limits of PB and equitably provisioning social development should be the point of concern in forthcoming times.

There are a few novelties of this work: First, it offers a pictorial portrait of the dynamic state of a set of socio-ecological indicators related to national priorities and scenarios in India. Our trend graphs show progress or regress over time that assists decisionmaking, and the amalgamation of environmental and social dimensions, along with the underlying role of economics, emphasize the triple bottle line feature of the sustainable development. Second, this work interconnects a multifaceted set of indicators in a relatively modest way, recognizes the gap in the underlying knowledge-base, and promotes new queries towards the elimination of social deprivation and achievement of environmental sustainability in India. Third, it provides India’s proximity to its environmental boundaries and its adequate level of social well-being. Fourth, the SDGs are supposed to be “action-oriented, concise and easy to communicate, limited in number, aspirational, global in nature and universally applicable to all countries, while taking into account different national realities, capacities and levels of development and respecting national policies and priorities” (Rio+20 outcome document, 2012). Almost all of these criteria have been met in this framework. Fifth, this work maintains a balance between simple and concise yet comprehensive, so that progress in all the SDGs in India can be understood (at least through one indicator for each SDG). Sixth, this study conveys insights into the challenges and complexities to develop appropriate indicators and boundaries at a national scale in India and focuses on zones where further research is needed to improvise this framework.

## Acknowledgement

We would like to thank Sk. Rohan Tanvir, The Institution of Engineers (India) for his assistance during the preparation of diagrams and Bishal Ghosh, Department of Economics, Presidency University for valuable suggestions.

---

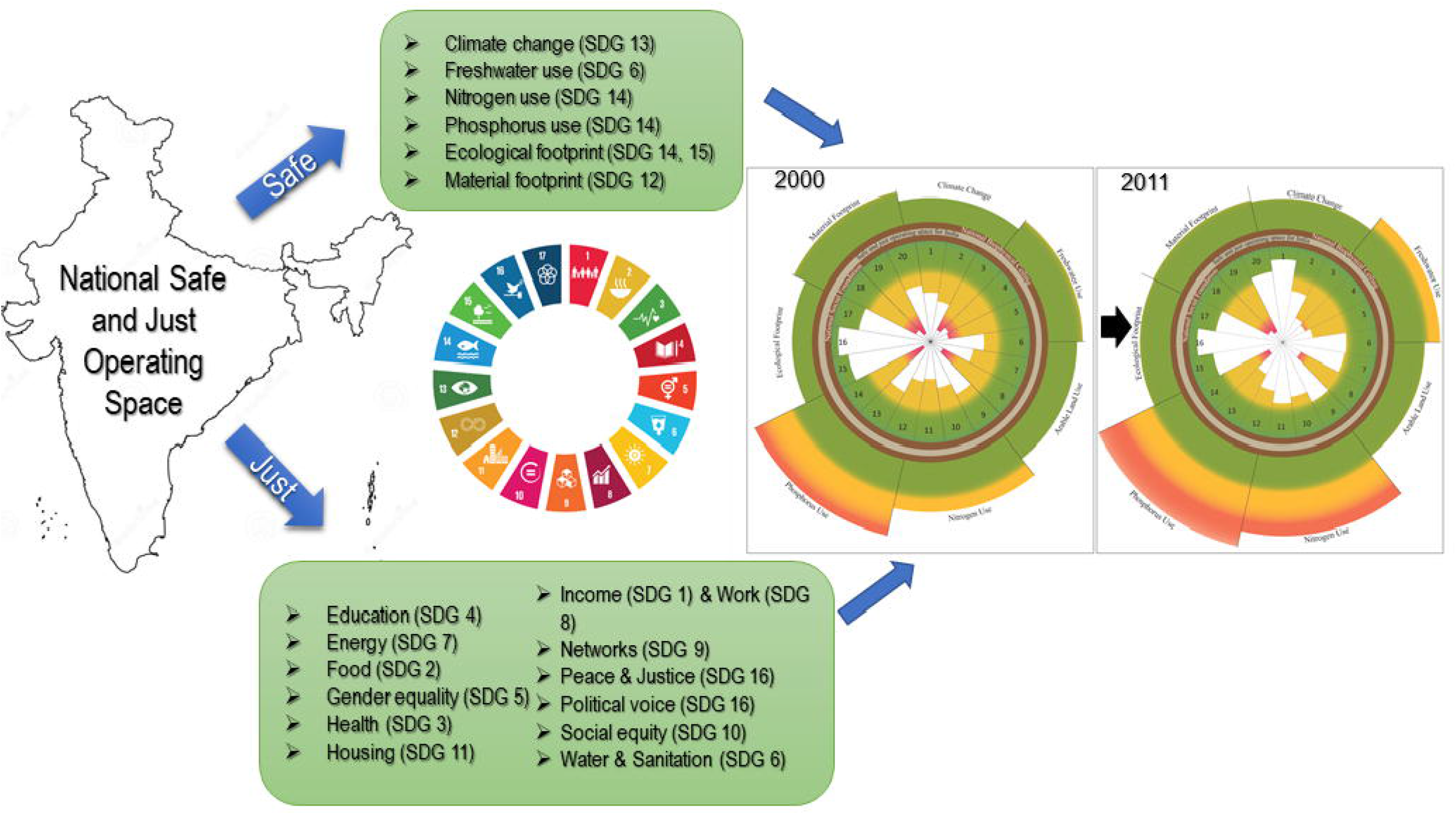

